# A functional genomic screen in *Saccharomyces cerevisiae* reveals divergent mechanisms of resistance to different alkylphosphocholine chemotherapeutic agents

**DOI:** 10.1101/2020.10.16.343244

**Authors:** Jacquelin M. Garcia, Michael J. Schwabe, Dennis R. Voelker, Wayne R. Riekhof

**Affiliations:** School of Biological Sciences, University of Nebraska – Lincoln, Lincoln, NE, USA; Department of Medicine, National Jewish Health, Denver, CO, USA; Division of Biology and Biomedical Sciences, Washington University, St. Louis, MO, USA; Department of Surgery, Creighton University School of Medicine, Omaha, NE, USA

## Abstract

The alkylphosphocholine (APC) class of antineoplastic and antiprotozoal drugs, such as edelfosine and miltefosine, are structural mimics of lyso-phosphatidylcholine (lyso-PC), and are inhibitory to the yeast *Saccharomyces cerevisiae* at low micromolar concentrations. Cytotoxic effects related to inhibition of phospholipid synthesis, induction of an unfolded protein response, inhibition of oxidative phosphorylation, and disruption of lipid rafts have been attributed to members of this drug class, however the molecular mechanisms of action of these drugs remain incompletely understood. Cytostatic and cytotoxic effects of the alkylphosphocholines exhibit variability with regard to chemical structure, leading to differences in effectiveness against different organisms or cell types. We now report the comprehensive identification of *Saccharomyces cerevisiae* titratable-essential gene and haploid non-essential gene deletion mutants that are resistant to the APC drug miltefosine (hexadecyl-*O*-phosphocholine). 58 strains out of ~5600 tested displayed robust and reproducible resistance to miltefosine. This gene set was heavily enriched in functions associated with vesicular transport steps, especially those involving endocytosis and retrograde transport of endosome derived vesicles to the Golgi or vacuole, suggesting a role for these trafficking pathways in transport of miltefosine to potential sites of action in the endoplasmic reticulum (ER) and mitochondrion. In addition, we identified mutants with defects in phosphatidylinositol-4-phosphate synthesis (TetO::*STT4*) and hydrolysis (*sac1*Δ), an oxysterol binding protein homolog (*osh2*Δ), a number of ER resident proteins, and multiple components of the eisosome. These findings suggest that ER-plasma membrane contact sites and retrograde vesicle transport are involved in the interorganelle transport of lyso-PtdCho and related lyso-phospholipid-like analogs to their intracellular sites of cytotoxic activity.

## Introduction

The alkylphosphocholine (APC) class of drugs are structural analogs of lyso-phosphatidylcholine (lyso-PtdCho), and have been extensively investigated as antineoplastic agents (C. Gajate et al., 2012; Consuelo Gajate & Mollinedo, 2014). They also act as effective anti-protozoal compounds, with potent activity against *Leishmania spp.* and other apicomplexan parasites (Saraiva *et al.* 2002; Santa-Rita *et al.* 2004; Verma *et al.* 2007; Aichelburg *et al.* 2008; Machado *et al.* 2010). *L. donovanii* has been investigated with regard to genetic mechanisms leading to drug resistance, and miltefosine-resistant strains have been identified in which a P_4_-ATPase (lipid flippase) at the plasma membrane is defective, and thus the resistant strain fails to import the toxic compound (Pérez-Victoria *et al.* 2006; Seifert *et al.* 2007; Weingärtner *et al.* 2010). The APC’s are also active against fungal pathogens such as *Candida albicans and Cryptococcus neoformans (Widmer et al. 2006; Vila et al. 2015)*, and the yeast *Saccharomyces cerevisiae* has served as a model for studies on drug resistance and elucidation of the molecular mechanism(s) of action of these drugs (Hanson *et al.* 2003; Cuesta-Marbán *et al.* 2013; Czyz *et al.* 2013). Although the body of work on APC resistance is extensive, particularly on mechanisms of action of edelfosine and miltefosine (Fig.1A), an understanding of the precise mechanism(s) of action of the APC’s remains incomplete, and the intracellular location of targets and mechanisms of action of these drugs appear to vary from compound to compound and between organisms and cell types.

**Figure 1:**
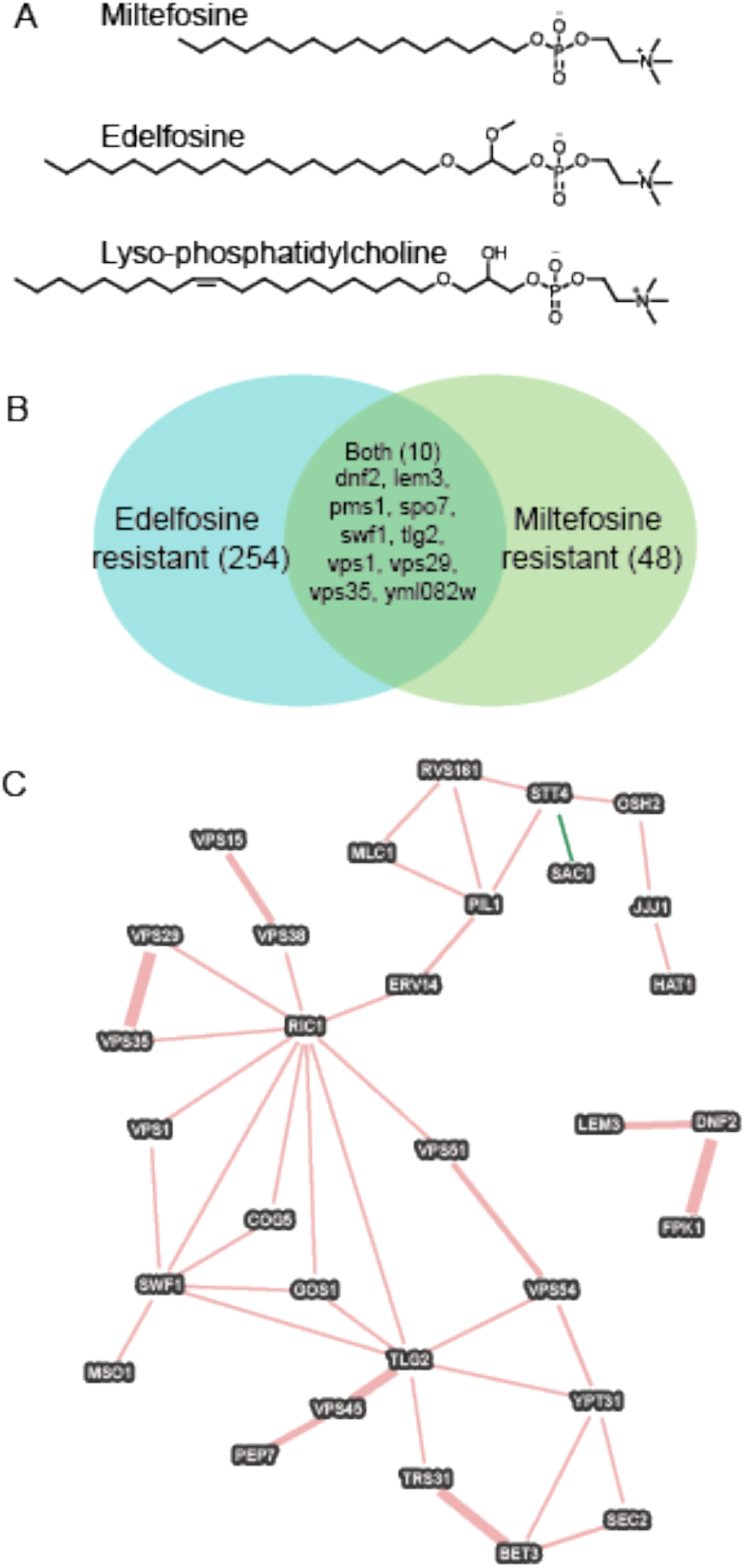
Overlap between previous and current screens for APC resistance. A, Structures of alkylphosphocholine compounds discussed in the text in comparison with lyso-Phosphatidylcholine.; B, Venn diagram illustrating the limited overlap in gene sets between the current study and previous studies on edelfosine resistance.; C, Physical and genetic interaction network of genes identified as being important for both miltefosine and edelfosine resistance, with additional network neighbors from the miltefosine-only screen described in this work.

A main mechanism of drug resistance, however, is clearly attributable to the loss of active transport of the APC’s into cells, which occurs via P_4_-family ATPase mediated internalization at the plasma membrane (Hanson *et al.* 2003; Riekhof and Voelker 2009). In the extensively studied *L. donovanii* system, disruption of the miltefosine transporter, LdMT (Pérez-Victoria *et al.* 2006; Seifert *et al.* 2007) or its non-catalytic β-subunit, LdRos3 (Pérez-Victoria *et al.* 2006) leads to drug resistance, similar to loss of the homologous *S. cerevisiae* proteins Dnf2p or Lem3p, respectively, which leads to edelfosine and miltefosine resistance in yeast (Hanson *et al.* 2003). A whole genome non-essential gene knockout screen for edelfosine resistant and hypersensitive mutants of *S. cerevisiae* was reported in a pair of studies (Cuesta-Marbán *et al.* 2013; Czyz *et al.* 2013). This work showed that edelfosine and miltefosine treatment causes loss of the proton efflux pump Pma1p from detergent resistant subdomains of the plasma membrane. This leads to an apparent mislocalization of Pma1p in APC treated cells, a corresponding defect in regulation of intracellular pH, and increased sensitivity to low extracellular pH. The edelfosine-resistant gene deletion mutant set these studies identified was enriched in genes encoding proteins with roles in endocytosis and endosomal transport, as well as in intracellular pH regulation.

We now report the comprehensive identification of miltefosine-resistant *S. cerevisiae* haploid deletion and titratable-essential gene strains. While there is overlap with the edelfosine-resistant mutant strains, we identified a substantial number of new mutant strains as miltefosine-resistant, suggesting that the mechanism(s) by which these compounds exert their cytotoxic effects are different, and that the mechanisms of fungal resistance to these drugs is not conserved across this drug class. Additionally, we identified an oxysterol-binding protein homolog (Osh2p) as being necessary for miltefosine sensitivity, and suggest that its presence at membrane contact sites is necessary for efficient APC dissemination to sites of action within the cell. We further show that proper Osh2p localization is likely to be dependent on the conent and localization of phosphatidylinositol-4-phosphate, as judged by mislocalization of Osh2p-GFP in TetO::*STT4* and *sac1*Δ strains.

## Materials and Methods

### Strains and growth conditions

All media components were from Fisher Scientific or Sigma-Aldrich. Routine growth and screening was conducted on YPD (1% w/v yeast extract, 2% w/v peptone, 2% w/v glucose) or YPGro (2% v/v glycerol instead of glucose) solidified with 1.5% w/v agar. Miltefosine was prepared as a 10 mg/ml stock solution in water, filter sterilized, and stored in frozen aliquots until just prior to use. Doxycycline was prepared as a 10 mM stock solution in ethanol and added to media (10 μM final concentration) as needed.

The *MAT*α deletion collection, constructed in parental strain BY4742 (Baker-Brachmann *et al.* 1998), and the tetracycline-repressible essential gene collection, constructed in strain R1158 (Mnaimneh *et al.* 2004), were purchased from Invitrogen. Additional *osh1*Δ, *osh2*Δ, *osh3*Δ, and *osh1*Δ *osh2*Δ *osh3*Δ strains in the SEY6210 background were provided by Tim Levine (University College London.) Initial screens were conducted by thawing 96-well glycerol stock plates, mixing with a stainless steel 96-pin tool (Dan-Kar model MC-96; Fisher Scientific), and dilution into 150 μl of sterile YPD medium in a 96-well plate. Approximately 3 μl of culture is transferred by each pin under these conditions, giving a ~50 fold dilution in the recipient plate. The diluted cultures were then pinned to solid YPD medium with or without 4 μg/ml (9.8 μM) miltefosine (Avanti Polar Lipids, Alabaster, AL, USA), incubated at 30 °C for up to 7 days, and visually monitored daily for growth of miltefosine-resistant patches. Cultures from the essential doxycycline-repressible (“Tet-off”) promoter collection (Mnaimneh *et al.* 2004) were screened similarly, except that an additional series of plates were used for the screen, which included 10 μg/ml doxycycline to effect repression of the essential library gene, as previously described (Wishart *et al.* 2005).

Miltefosine-resistant strains were identified and subcultured for further studies. Resistant strains were colony-purified from patches on the uninhibited miltefosine-free YPD replica plates, and single colonies were picked and grown to saturation in YPD. A 10-fold dilution series was prepared in 96-well plates and pinned to YPD (4 μg/ml miltefosine) or YPGro (1 μg or 4 μg/ml miltefosine) and growth assessed after 48 or 96 h, respectively. Strains that showed reproducible miltefosine resistance were thus identified, and categorized based on degree of sensitivity with glucose or glycerol as carbon sources.

### Bioinformatic analyses

Primary and secondary screening led to identification of 58 genes which, when deleted or repressed, led to a reproducible miltefosine-resistant growth phenotype. Gene ontology analysis was performed with YeastMine tools available at the Saccharomyces Genome Database website (www.yeastgenome.org), and gene set analysis was conducted with YeastNet version 3.0 at www.inetbio.org/yeastnet (Kim *et al.* 2014).

### Microscopy

Strain BY4742 and the isogenic *sac1*Δ::*KanMX* deletion strain were transformed to uracil prototrohy with plasmid pTS312, a *URA3 CEN* plasmid expressing a C-terminal GFP fusion of Osh2p, which was a gift of Christopher Beh (Simon Fraser University.) Cultures were grown overnight in SC -Ura and imaged on an Evos FL inverted microscope with GFP fluorescence cube.

## Results and Discussion

### Screening for miltefosine resistant strains identifies a network of 58 highly connected genes enriched in membrane and trafficking functions

Table 1 provides a list of the genes that were identified as showing reproducible resistance to 4 μg/ml miltefosine on YPD, and that rescreened as positive after colony purification of original stocks and assessment with a 5-fold serial dilution spot-test. Relevant molecular structures are shown in Fig, 1A. Fig. 2 shows a typical re-screening result for a subset of 6 mutants from the haploid MATα BY4742 based mutant collection. These initial results demonstrated that the miltefosine-resistance phenotype is tractable in the context of a genome-wide deletion screen conducted on solid medium. Figure 5 shows a result typical of the essential-gene Tet-off screening phenotype (TetO::*STT4*), in which we selected strains that showed robust growth in the presence of miltefosine when doxycycline was present (gene repressed), and weak or absent growth in its absence (gene expressed normally)

**Table 1:** List of genes identified in the miltefosine-resistance screen. Genes are listed in order of chromosomal position as noted from the standard SGD identifier, with gene name (if available), assessment of whether the mutant had previously been identified as resistant to edelfosine or miltefosine, and a description of the gene product.

| Gene ID | Name | Previous ID? | Description |
| --- | --- | --- | --- |
| YAL009W | SPO7 | Yes | Subunit of Nem1p-Spo7p phosphatase holoenzyme |
| YAL039C | CYC3 | No | Cytochrome c heme lyase |
| YBR097W | VPS15 | No | Protein kinase involved in vacuolar protein sorting |
| YBR184W |  | No | Putative protein of unknown function |
| YCR009C | RVS161 | No | Amphiphysin-like lipid raft protein |
| YCR067C | SED4 | No | ER-protein that stimulates Sar1p GTPase activity |
| YDL019C | OSH2 | No | Oxysterol-binding protein homolog family member |
| YDR027C | VPS54 | No | Golgi-associated retrograde protein complex |
| YDR028C | REG1 | No | Subunit of type 1 protein phosphatase Glc7p |
| YDR093W | DNF2 | Yes | Aminophospholipid translocase (flippase) |
| YDR095C |  | No | Dubious ORF overlapping *DNF2* |
| YDR097C | MSH6 | No | Required for mismatch repair in mitosis and meiosis |
| YDR126W | SWF1 | Yes | Palmitoyltransferase of SNARE proteins |
| YDR323C | PEP7 | No | Vesicle-mediated vacuolar protein sorting |
| YDR472W | TRS31 | No | Transport protein particle (TRAPP) I-III |
| YER031C | YPT31 | No | Rab family GTPase |
| YFL047W | RGD2 | No | GTPase-activating protein for Cdc42p and Rho5p |
| YGL012W | ERG4 | No | C-24(28) sterol reductase |
| YGL054C | ERV14 | No | COPII-coated vesicle protein |
| YGL095C | VPS45 | No | Protein of the Sec1p/Munc-18 family |
| YGL106W | MLC1 | No | Essential light chain for Myo1p |
| YGL158W | RCK1 | No | Protein kinase involved in oxidative stress response |
| YGR086C | PIL1 | No | Eisosome core component |
| YGR141W | VPS62 | No | Vacuolar protein sorting (VPS) protein |
| YHL028W | WSC4 | No | Endoplasmic reticulum (ER) membrane protein |
| YHL031C | GOS1 | No | v-SNARE protein involved in Golgi transport |
| YHR012W | VPS29 | Yes | Subunit of retromer complex |
| YHR108W | GGA2 | No | Regulates Arf1p, Arf2p to facilitate Golgi trafficking |
| YJL154C | VPS35 | Yes | Endosomal subunit of retromer complex |
| YKL212W | SAC1 | No | PtdIns-4-phosphate phosphatase |
| YKR001C | VPS1 | Yes | Dynammin-like GTPase required for vacuolar sorting |
| YKR019C | IRS4 | No | EH domain-containing protein |
| YKR020W | VPS51 | No | Golgi-associated retrograde protein complex |
| YKR068C | BET3 | No | Transport protein particle (TRAPP) complexes I-III |
| YLR039C | RIC1 | No | Retrograde transport to the cis-Golgi network |
| YLR082C | SRL2 | No | Protein of unknown function |
| YLR093C | NYV1 | No | v-SNARE component of vacuolar membrane |
| YLR305C | STT4 | No | Phosphatidylinositol-4-kinase |
| YLR360W | VPS38 | No | Vps34p phosphatidylinositol-3-kinase complex |
| YML052W | SUR7 | No | Plasma membrane component of eisosomes |
| YML082W |  | Yes | Protein of unknown function |
| YMR032W | HOF1 | No | Regulates actin cytoskeleton organization |
| YNL051W | COG5 | No | Conserved oligomeric Golgi complex subunit |
| YNL058C |  | No | Putative protein of unknown function |
| YNL082W | PMS1 | Yes | ATP-binding protein required for mismatch repair |
| YNL227C | JJJ1 | No | Stimulates the ATPase activity of Ssa1p |
| YNL272C | SEC2 | No | Guanyl-exchange factor for small G-protein Sec4p |
| YNL323W | LEM3 | Yes | Beta-subunit of Lem3p flippase |
| YNR047W | FPK1 | No | Ser/Thr protein kinase activating Dnf2p |
| YNR049C | MSO1 | No | Lipid-interacting protein in SNARE assembly |
| YOL018C | TLG2 | Yes | Syntaxin-like t-SNARE |
| YOL107W |  | No | Putative protein of unknown function |
| YOR311C | DGK1 | No | Diacylglycerol kinase |
| YPL001W | HAT1 | No | Hat1p-Hat2p histone acetyltransferase complex |
| YPL028W | ERG10 | No | Acetyl-CoA C-acetyltransferase |
| YPR089W |  | No | Protein of unknown function |
| YPR117W |  | No | Putative protein of unknown function |
| YPR151C | SUE1 | No | Degradation of unstable forms of cytochrome c |

**Figure 2:**
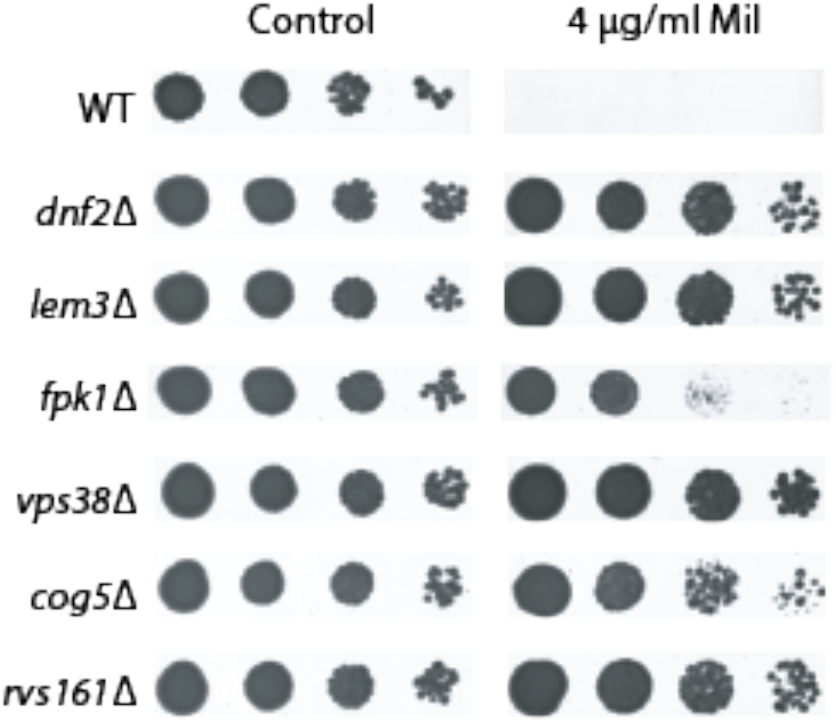
Miltefosine-resistance phenotypes. Primary screening led to the identification of strains with a range of resistance phenotypes. Strains were grown overnight in YPD and a 5-fold serial dilution series was prepared and pinned to solid YPD agar with or without miltefosine. Plates were photographed after 3 days. The densest spots correspond to the 1:25 dilution, and lightest correspond to 1:3125 dilution. A range of phenotypes from weakly (fpk1Δ) to modestly (cog5Δ) to strongly (dnf2Δ, vps38Δ) resistant are noted.

When screening mutant libraries for strains with alterations in a phenotype of interest, it is useful to assess whether the screen has identified mutants in a subset of genes with related functions, or whether the screen has identified a set of diverse genes with little in common with regard to function or localization. We approached this question by assessing the connectivity of the identified gene set with regard to systems-based epistasis screens, protein-protein interaction studies, and other measures, such as shared protein domain architectures and co-citation indices. We also analyzed the enrichment of gene ontology (GO) terms associated with the gene set. The method of assessment of connectivity makes use of receiver operating characteristic (ROC) curve analysis (McGary *et al.* 2007), in which the area under the ROC curve is assessed in relation to that expected for a randomly chosen subset of genes. The results of ROC analysis for our set of 58 genes is given in Fig. 3. This approach makes use of the YeastNet v3 database (Kim *et al.* 2014) to assess whether a set of genes are more connected to each other than they are to a random subset of the genome. This approach led us to discover that miltefosine resistance is a powerful selection for mutants in a subset of genes that are highly connected via genetic interactions, protein:protein interactions, and co-citation indices with one another, with a p-value of 8.3 × 10^−11^. This means that that the chances of identifying a random set of genes with this level of connectivity are on the order of 1 in 10^−10^. As a control, we selected two random pools of 58 genes each and analyzed their connectivity by ROC analysis as for the miltefosine-resistance gene set. As expected, these subsets of genes displayed no significant numbers of connections beyond that which would be expected from random chance.

**Figure 3:**
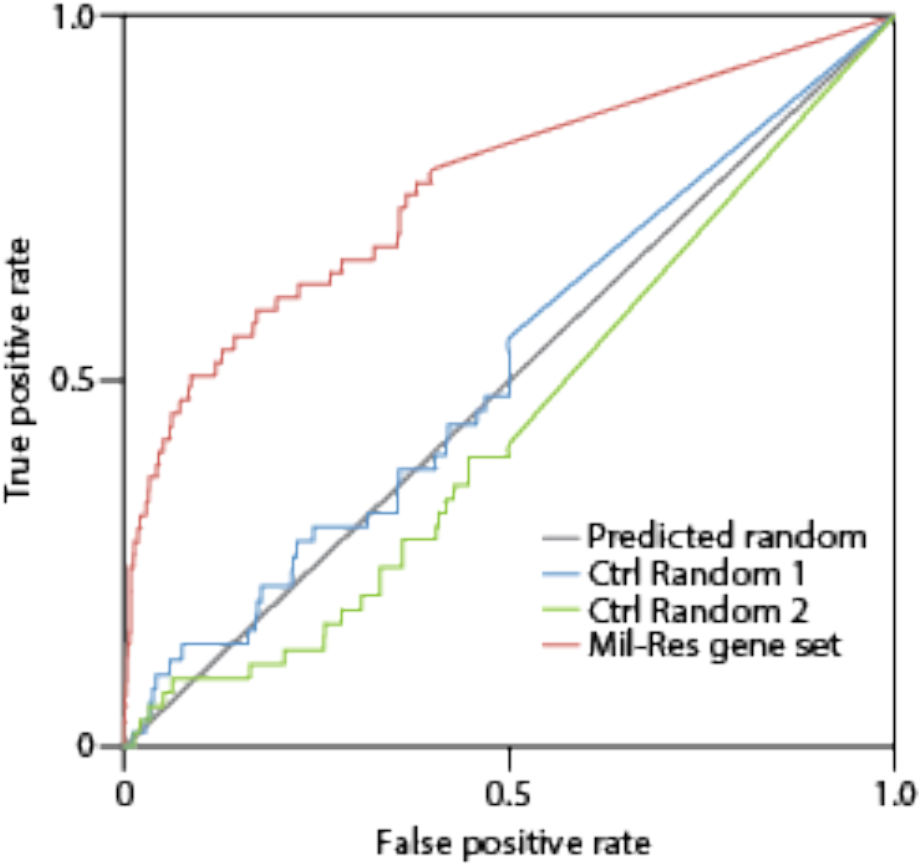
Receiver-operating characteristic curve analysis. The gene set of reproducibly miltefosine resistant mutants was analyzed with gene-set enrichment analysis tools from the YeastNet v3 package at https://www.inetbio.org/yeastnet/ as described in the text. Two sets of randomly selected genes were analysed for comparison, and the miltefosine-resistant gene set was found to identify genes that were significantly connected to each other via protein:protein interactions, genetic epistasis, shared domain architectures, and co-citation indices with an E value of less than 1 × 10^−10^.

A second method of gene set analysis was performed by assessing the enrichment of gene ontology (GO) terms as a measure of relatedness, and allowed us to identify molecular components and processes that are significantly over-represented in the miltefosine-resistant mutant collection. Table 2 provides a list of GO terms that are over-represented in the miltefosine-resistant gene set. As might be expected for a set of mutants with resistance to a membrane-perturbing agent, GO components and processes associated with membrane assembly, protein trafficking, and transport through the endomembrane system are highly enriched in this analysis.

**Table 2:** Gene-ontology term enrichment. Genes identified in the mutant screen were analyzed with Gene Ontology (GO) based gene-set enrichment analysis tools available at www.yeastgenome.org. p-values were calculated based on the hypergeometric test, and GO terms with p < 0.005 are included in the table.

| GO identifier | GO Component (C) or Process (P) | Total for<br>term | Identified | Fold<br>enrichment | p-value |
| --- | --- | --- | --- | --- | --- |
| GO:0005794 | C:Golgi apparatus | 214 | 17 | 8.2 | 6.92E-12 |
| GO:0000139 | C:Golgi membrane | 121 | 10 | 8.5 | 1.77E-07 |
| GO:0031201 | C:SNARE complex | 25 | 5 | 20.7 | 3.26E-06 |
| GO:0005768 | C:endosome | 114 | 8 | 7.2 | 1.16E-05 |
| GO:0016020 | C:membrane | 1718 | 31 | 1.9 | 5.54E-05 |
| GO:0010008 | C:endosome membrane | 59 | 5 | 8.8 | 0.0002392 |
| GO:0000938 | C:GARP complex | 4 | 2 | 51.6 | 0.0005458 |
| GO:0034272 | C:phosphatidylinositol 3-kinase<br>complex II | 4 | 2 | 51.6 | 0.0005458 |
| GO:0030906 | C:retromer complex, inner shell | 4 | 2 | 51.6 | 0.0005458 |
| GO:0030904 | C:retromer complex | 5 | 2 | 41.3 | 0.0009041 |
| GO:0006810 | P:transport | 793 | 22 | 2.9 | 1.74E-06 |
| GO:0015031 | P:protein transport | 389 | 17 | 4.5 | 7.43E-08 |
| GO:0016192 | P:vesicle-mediated transport | 140 | 10 | 7.4 | 6.98E-07 |
| GO:0006897 | P:endocytosis | 88 | 8 | 9.4 | 1.64E-06 |
| GO:0006896 | P:Golgi to vacuole transport | 24 | 5 | 21.5 | 2.63E-06 |
| GO:0006623 | P:protein targeting to vacuole | 48 | 5 | 10.8 | 8.87E-05 |
| GO:0042147 | P:retrograde transport, endosome to<br>Golgi | 19 | 5 | 27.2 | 7.46E-07 |
| GO:0006914 | P:autophagy | 51 | 4 | 8.1 | 0.001407 |
| GO:0032258 | P:CVT pathway | 38 | 4 | 10.9 | 0.0004564 |
| GO:0006906 | P:vesicle fusion | 20 | 4 | 20.7 | 3.41E-05 |
| GO:0006895 | P:Golgi to endosome transport | 14 | 3 | 22.1 | 0.0002905 |
| GO:0006869 | P:lipid transport | 21 | 3 | 14.8 | 0.001012 |
| GO:0045053 | P:protein retention in Golgi apparatus | 11 | 3 | 28.2 | 0.0001344 |
| GO:0000011 | P:vacuole inheritance | 18 | 3 | 17.2 | 0.0006335 |
| GO:0006904 | P:vesicle docking involved in<br>exocytosis | 10 | 3 | 31.0 | 9.84E-05 |
| GO:0048017 | P:inositol lipid-mediated signaling | 5 | 2 | 41.3 | 0.0009041 |
| GO:0060988 | P:lipid tube assembly | 3 | 2 | 68.9 | 0.0002746 |

### The miltefosine resistant gene deletion set shows little overlap with a previously described edelfosine resistance screen

As noted in Figure 1, approximately 264 deletion mutant strains were previously identified as showing some degree of edelfosine resistance (Cuesta-Marbán *et al.* 2013; Czyz *et al.* 2013). Our current study identified 58 mutants showing resistance to miltefosine, and of those, only 10 mutants were shared between the edelfosine and miltefosine resistance sets, although this may be a slight underestimate given that the edelfosine resistance screen did not encompass the essential gene titratable-promoter collection. Figure 1C shows a composite genetic and physical interaction map of edelfosine-miltefosine resistant shared genes and their directly interacting partners in the miltefosine resistance set, and identifies the t-SNARE TLG2 as a key hub in the interaction diagram for the shared edelfosine-miltefosine resistant gene set.

A *tlg2*Δ strain was identified along with *dnf2*Δ, *lem3*Δ, and other components of endosome-plasma membrane recycling in a screen for strains resistant to the lantibiotic peptide Ro 09-0198 (Takagi *et al.* 2012). This work also showed that an EGFP-tagged Dnf2p reporter was mislocalized in the *tlg2*Δ background, suggesting that the Ro-0198 sensitive phenotype of *tlg2*Δ is due to mislocalization of Dnf2p from the plasma membrane to endosomes. Disruption or mislocalization of this lipid flippase would thus result in the accumulation of phosphatidylethanolamine on the outer leaflet of the plasma membrane, which is the ligand for this cytolytic peptide. Taken together with our results, this suggests that localization and proper function of the Dnf2p-Lem3p complex is dependent on Tlg2p and a small cadre of proteins to which it is functionally linked, and that the genes in the intersection of the Venn diagram of Fig. 1B and in the interaction diagram of Fig. 1C encompass the core components necessary for flippase function, activity, and localization at the plasma membrane. We thus propose that while the core components of lysophospholipid uptake (the flippase core components and factors regulating its proper localization and function) are broadly conserved as determinants of APC sensitivity, individual members of this this drug class exert their cytotoxic effects by interacting with multiple and variable intracellular targets after their import via the flippase.

### Disruption of phosphatidylinositol-4-phosphate homeostasis and the oxysterol-binding protein homolog Osh2p alters miltefosine sensitivity

The proteins disrupted in a subset of functionally related miltefosine-resistant mutants (*osh2*Δ, *sac1*Δ, and TetO::*STT4*) are involved in the function and localization of the oxysterol binding protein homolog Osh2p. Osh2p contains an oxysterol-binding domain at its C-terminus as well as several protein-protein and protein:lipid interaction motifs at its N-terminus. These interaction motifs include Anykyrin-repeats which are likely to interact with other, currently unidentified, proteins (Beh *et al.* 2001), a Pleckstrin-homology (PH) domain that interacts with PtdIns-4-P (Roy and Levine 2004), and a “two phenylalanines in an acidic tract” (FFAT) motif which interacts with the ER resident proteins Scs2p and Scs22p (Loewen *et al.* 2003; Kaiser *et al.* 2005; Loewen and Levine 2005).

The phosphatidylinositol (PtdIns) 4-kinase Stt4p (Yoshida *et al.* 1994; Cutler *et al.* 1997) generates PtdIns-4-P at the plasma membrane (Foti *et al.* 2001; Baird *et al.* 2008), and is involved in actin polymerization and endocytosis, as well as in transport of PtdSer from the ER to the site of Psd2p (Trotter *et al.* 1998). The pool of PtdIns-4-P generated by Stt4p is degraded by the phosphoinositide phosphatase Sac1p (Foti *et al.* 2001), which has additional functions in lipid trafficking and metabolism (Guo *et al.* 1999; Rivas *et al.* 1999; Hughes *et al.* 2000; Tahirovic *et al.* 2005; Riekhof *et al.* 2014; Tani and Kuge 2014). The strains *sac1*Δ, *osh2*Δ, and TetO::*STT4* were all identified as miltefosine resistant (Figs. 4 and 5), suggesting that the PtdIns-4-P cycle governed by Stt4p and Sac1p might be involved in the proper localization of Osh2p, and that mislocalization of Osh2p might lead to a defect in miltefosine uptake and distribution to intracellular targets. This idea was tested by expressing a GFP-tagged form of Osh2p in a wild-type and *sac1*Δ background. As shown in Fig. 6, Osh2p in the wild type is localized in a punctate pattern at the cell periphery, while in the *sac1*Δ mutant Osh2p is localized to intracellular structures and absent from the cell periphery.

**Figure 4:**
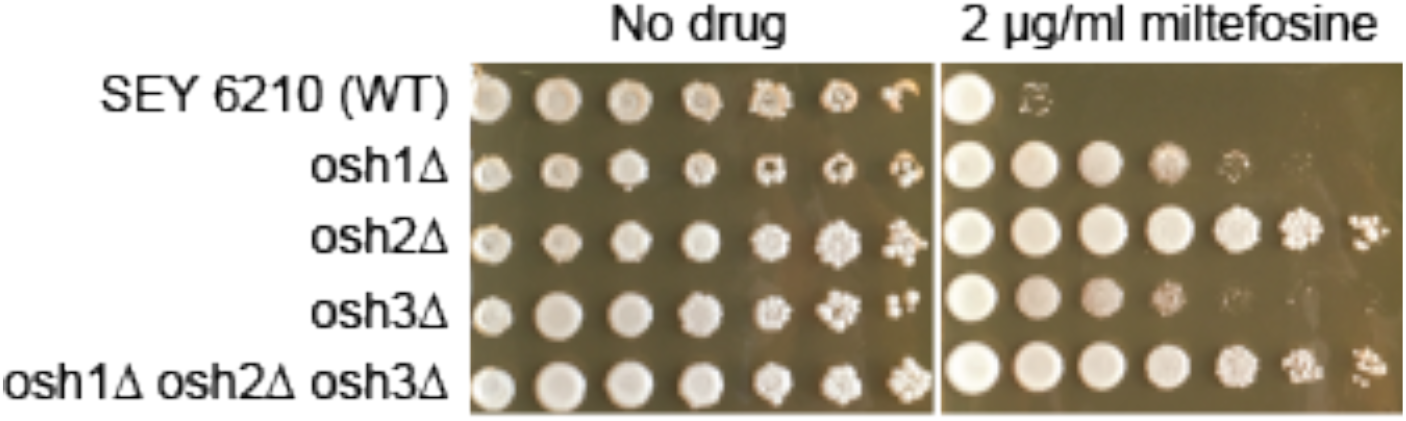
Oxysterol binding protein homologs are necessary for wild-type miltefosine sensitivity. An osh2Δ strain was identified in the primary screen conducted in the BY4742 background. To confirm the role of Osh2p and its orthologs Osh1p and Osh3p and determine whether there were background specific differences in sensitivity, we assessed the growth of strain SEY6210 and isogenic *osh1*Δ, *osh2*Δ, *osh2*Δ, and the triple mutant. Overnight cultures were subjected to 5-fold serial dilution and pinned to solid media as described in the text.

**Figure 5:**
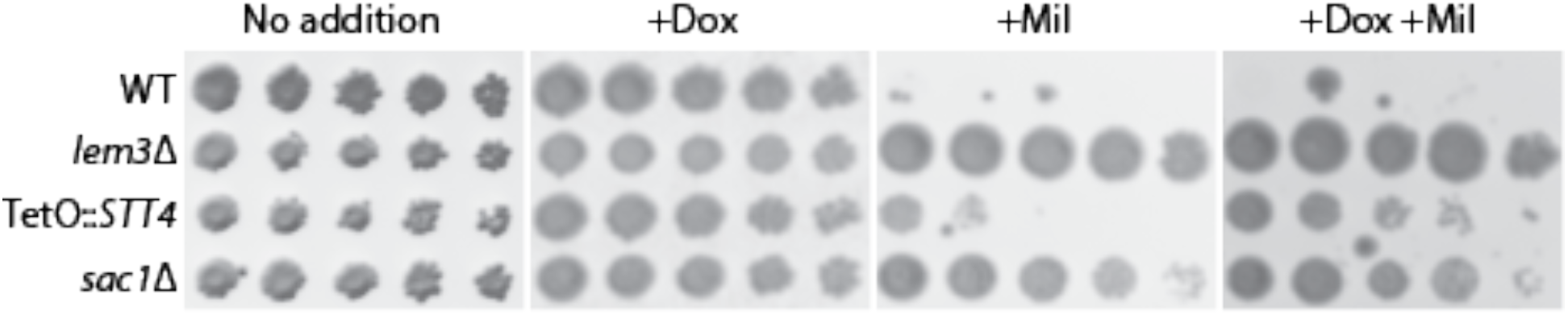
The phosphatidylinositol-4-kinase/phosphatase cycle is a determinant of miltefosine sensitivity. The essential gene *STT4,* encoding the major PtdIns-4-kinase isoform in yeast, was identified in the essential-titratable gene collection. Inhibition of *STT4* expression by including doxycycline in the growth media (10 μM) led to mitefosine resistance (4 μg/ml.) The PtdIns-4-P phosphatase Sac1p, was also identified and a *sac1*Δ strain was included in this series, and *lem3*Δ was included as a previously characterized miltefosine resistant positive control. 5-fold serial dilutions were prepared and pinned to solid media as described in the text.

**Figure 6:**
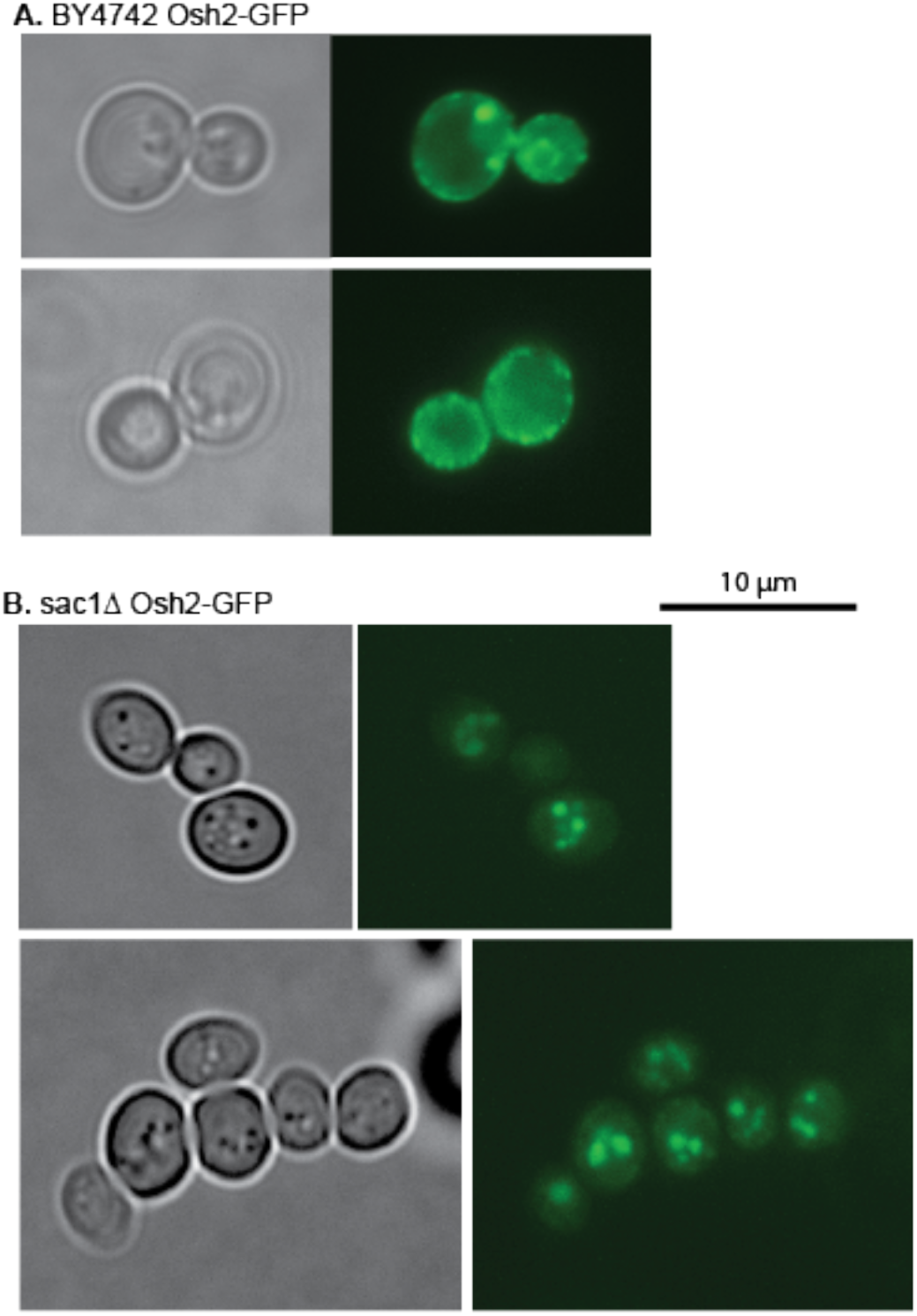
Osh2p is mislocalized in a sac1Δ mutant background. Osh2-GFP expressed from a *URA3* CEN plasmid under its native promoter was introduced into the BY4742 wild type and isogenic sac1Δ mutant strains. Strains were grown in SC -Ura media overnight and images were acquired on an Evos-*Fl* inverted microscope with a GFP light cube using a 100x oil-immersion objective.

### Summary

Previous studies in *S. cerevisiae* (Cuesta-Marbán *et al.* 2013; Czyz *et al.* 2013) have identified a subset of genes that, when deleted, confer resistance to the APC analog edelfosine. While members of this drug class share superficial similarities to lysophosphatidylcholine in their structures, it has remained an open question as to the degree of overlap in their mechanisms of action. Our current work identifies the plasma membrane flippase Dnf2p and factors required for its correct localization and function as being shared between edelfosine, miltefosine and likely other members of this drug class with regard to the specific and active transport of the compounds into the cell. However, unlike the edelfosine studies, we did not identify factors that regulate intracellular pH or alter the function of the plasma membrane proton pump Pma1p as having a role in altered miltefosine sensitivity. We did however identify a new subset of genes as determinants of wild-type miltefosine sensitivity, and chose to assess the roles of the PtdIns-4-P cycle and Osh2 localization in more detail. One interpretation of this data is that after the initial “flip” across the plasma membrane by Dnf2p/Lem3, miltefosine is transported to one or more sites of cytotoxic activity by non-vesicular transport routes that are dependent on the proper localization of the oxysterol-binding protein homolog Osh2p. Future work on this and other clusters of genes identified in this screen will provide additional insight into the mechanism(s) of action of miltefosine and other members of the APC drug class.

## Acknowledgments

This work was supported by UNL Undergraduate Creative Activities and Research Enhancement (UNL-UCARE) and Ronald E. McNair Scholars program fellowships (to J.E.G.); National Science Foundation grant EPS-1004094 (to W.R.R.) and an Institutional Research Grant from the Fred and Pamela Buffett Cancer Center and the American Cancer Society (to W.R.R.) We thank Christopher Beh (Simon Fraser University, Vancouver, BC, Canada) and Tim Levine (University College London, London, UK) for providing yeast strains and plasmids as noted in the text.

## Author contributions

J.E.G. acquired funding, performed research, assisted with study design, performed data analysis, prepared figures, and provided feedback on the manuscript draft; M.J.S performed research, assisted with data analysis, and provided feedback on the manuscript draft; D.R.V. provided space and assistance for generating the conceptual framework that led to this study, and provided feedback on early stages of study design and on the manuscript draft; W.R.R. designed the study, acquired funding, performed and directed research, and wrote the manuscript.

